# Next-generation genetic sexing strain establishment in the agricultural pest *Ceratitis capitata*

**DOI:** 10.1101/2023.09.29.560088

**Authors:** Serafima Davydova, Junru Liu, Nikolay P. Kandul, W. Evan Braswell, Omar S. Akbari, Angela Meccariello

## Abstract

Tephritid fruit fly pests pose an increasing threat to the agricultural industry due to their global dispersion and a highly invasive nature. Here we showcase the feasibility of an early-detection SEPARATOR sex sorting approach through using the non-model Tephritid pest, *Ceratitis capitata*. This system relies on female-only fluorescent marker expression, accomplished through the use of a sex-specific intron of the highly-conserved *transformer* gene from *C. capitata* and *Anastrepha ludens*. The herein characterized strains have 100% desired phenotype outcomes, allowing accurate male-female separation during early development. Overall, we describe an antibiotic and temperature-independent sex-sorting system in *C. capitata*, which, moving forward, may be implemented in other non-model Tephritid pest species. This strategy can facilitate the establishment of genetic sexing systems with endogenous elements exclusively, which, on a wider scale, can improve pest population control strategies like sterile insect technique.

## Introduction

The Mediterranean fruit fly (medfly), *Ceratitis capitata* (*Tephritidae*), is among the most notorious and vigorous agricultural pests. It has a vast host range of over 260 fruit and vegetable species, and a worldwide distribution characterized by ongoing rapid expansions (1 – 4). Its invasive nature is nourished by factors like increased food trading fueled by the growing demand for produce (1) and climate change (3, 5 – 7). The medfly often acts as a model for other Tephritid pests such as the Mexican fruit fly, *Anastrepha ludens*, due to its easier laboratory-based rearing and a shorter life cycle.

Tephritid pests are targeted for population control through sterile insect technique (SIT) as part of area-wide integrated pest management (AW-IPM) programs implemented internationally (8). SIT relies on mass rearing, irradiation-based sterilization of flies and their sequential release into the wild (9, 10). The aim of this strategy is to promote mating of released sterile males with wild females, suppressing population growth (10). In general, both males and females have been released simultaneously in SIT (11). Male-only releases, however, were shown to be more cost-effective than co-releases with females (12). Additionally, male-only release schemes reduce crop damage invoked by egg laying and encourage wider sterile male dispersal allowing greater mating frequency with wild females (13, 14).

To eliminate the necessity for sex-sorting at adulthood in SIT and allow male-only releases, multiple cost-effective genetic sexing strains (GSSs) were developed in Tephritids (15 – 18). Traditionally, GSS requires a selective marker, for instance conditional female development arrest or puparium coloration which is linked to the Y chromosome through translocation (15). Such strains have been implemented in *C. capitata* SIT rearing (19); and developed in multiple other Tephritid species (16 – 18). The largest problem with Y-linked GSSs is the associated genetic instability caused by recombination and thus desired phenotype loss (15). Another complication with the currently used GSSs is the partial sterility of the males which creates rearing difficulties (20). To tackle this, novel scalable sexing strategies are required, boasting superior genetic stability and lower fitness costs (11, 21).

To address genetic instability of Y-linked GSSs, multiple Y-independent GSSs have been developed in fruit flies (22 – 26). The majority of Y-independent GSS approaches rely on the sex-specific alternative splicing properties of a sex determination gene, *transformer* (*tra)* (22, 24 – 26). Among Tephritids, the role of *tra* is best characterized in the medfly, although its functions are well-conserved across the family (27). The default mechanism of medfly embryonic development involves maternal and zygotic *C. capitata tra* (*Cctra*) and *C. capitata tra2* (*Cctra2*) which enable a positive feedback loop of female-specific splicing of *Cctra* (28 – 30). This female form of Cctra is needed for female-specific splicing of downstream *doublesex* and *fruitless* genes, conserved across the Tephritidae family (27, 31 – 35). In XY male embryos, the female-specific splicing of *tra* is blocked by Maleness on Y (*MoY*) factor, transcribed from the Y chromosome (36). In the presence of *MoY, tra* is spliced with stop codon-containing male-specific exons resulting in premature translation arrest and thus the lack of functional Cctra production (37). For GSS establishment, the *C. capitata tra* intron was used to induce female-specific lethality in two tetracycline-dependent systems in the medfly (22, 25) and similarly in *A. ludens* (26). Although highly efficient, these sexing strains require tetracycline supply for line maintenance which affects mitochondrial function and fitness of released males (38 – 41), as well as becoming costly for production on a mass scale (22, 25, 26). Hence alternative approaches would be beneficial to alleviate the reliance of Y-independent GSSs on tetracycline.

In this study, we showcase a sex-sorting method in *C. capitata* by exploiting Sexing Element Produced by Alternative RNA-splicing of a Transgenic Observable Reporter (SEPARATOR). This strategy was previously used in *Drosophila melanogaster* as a proof of principle, and further demonstrated in *Aedes aegypti* mosquitoes and enabled sex-sorting of early L_1_ is through large particle flow cytometry using a complex object parametric analysis and sorting (COPAS) instrument (42, 43). The transgenic SEPARATOR lines generated here harbor a cassette with one constituent, and one *tra* intron-containing fluorescence markers. Thus, phenotypically, the males and females can be easily distinguished through fluorescence screening from early development. We demonstrate the feasibility of such a system using both the endogenous *C. capitata* and *A. ludens tra* introns. Our work provides a basis for accessible cross-species transfer across Tephritid family members with a shorter research history, including *A. ludens*. Moreover, the strategy described here can serve as a cornerstone for seamless endogenous GSS generation through insertion of the *tra* intron into genes such as *white pupae* and *ebony* for their sex-specific expression (15, 18). Such an approach is feasible due to the recent successful implementation of homology-directed repair in the medfly (44, 45).

## Results

### Establishment and characterization of fluorescent SEPARATOR strains

To engineer the SEPARATOR genetic cassettes for *Ceratitis capitata*, we utilized the *transformer* (*tra*) intron of *C. capitata* or *A. ludens* to generate 795H1 or 795K1 construct, respectively. Importantly, both constructs included the *DsRed* coding sequence separated by the *tra* intron and expressed under the *Opie2* promoter, and a dominant *Hr5-IE1-eGFP* marker (Figure 1a). We used germline transformation to induce *piggyBac*-reliant integrations for 795H1 and 795K1 constructs (Supplementary Table 1). Two separate strains for each construct were successfully characterized through inverse PCR (H-001 and H-002 for 795H1; K-001 and K-002 for 795K1) (Supplementary Figure 1 and Supplementary Table 2). Homozygosity for all strains was achieved via sibling crosses at G_2_ thereafter. We noted no DsRed+/GFP+ males or DsRed-/GFP+ females for any of the 4 homozygous strains during routine screenings for 8 consecutive generations (from G_3_ to G_10_). This was indicative of sex-specific splicing of DsRed, and thus the success of *tra* intron selection for both *C. capitata* and *A. ludens*. For verification, we expanded the strains and characterized the entire populations at G_9_ and G_10_ (Figure 2a). As expected, no DsRed+/GFP+ males or DsRed-/GFP+ females were detected. Transgenic individuals with only two phenotypes: DsRed+/GFP+ females (48.8 %) and DsRed-/GFP+ males (51.2 %) were noted across the four strains (Supplementary Table 3). Chi-squared tests for observed and expected sex ratios showed no statistical significance for any of the four strains. Overall, 100% transgenic flies expressed the desired fluorescence phenotypes across both *C. capitata* (795H1) and *A. ludens* (795K1) *tra* intron-harboring medflies across G_9_ and G_10_ with a total of 3,787 flies counted.

**Figure 1:**
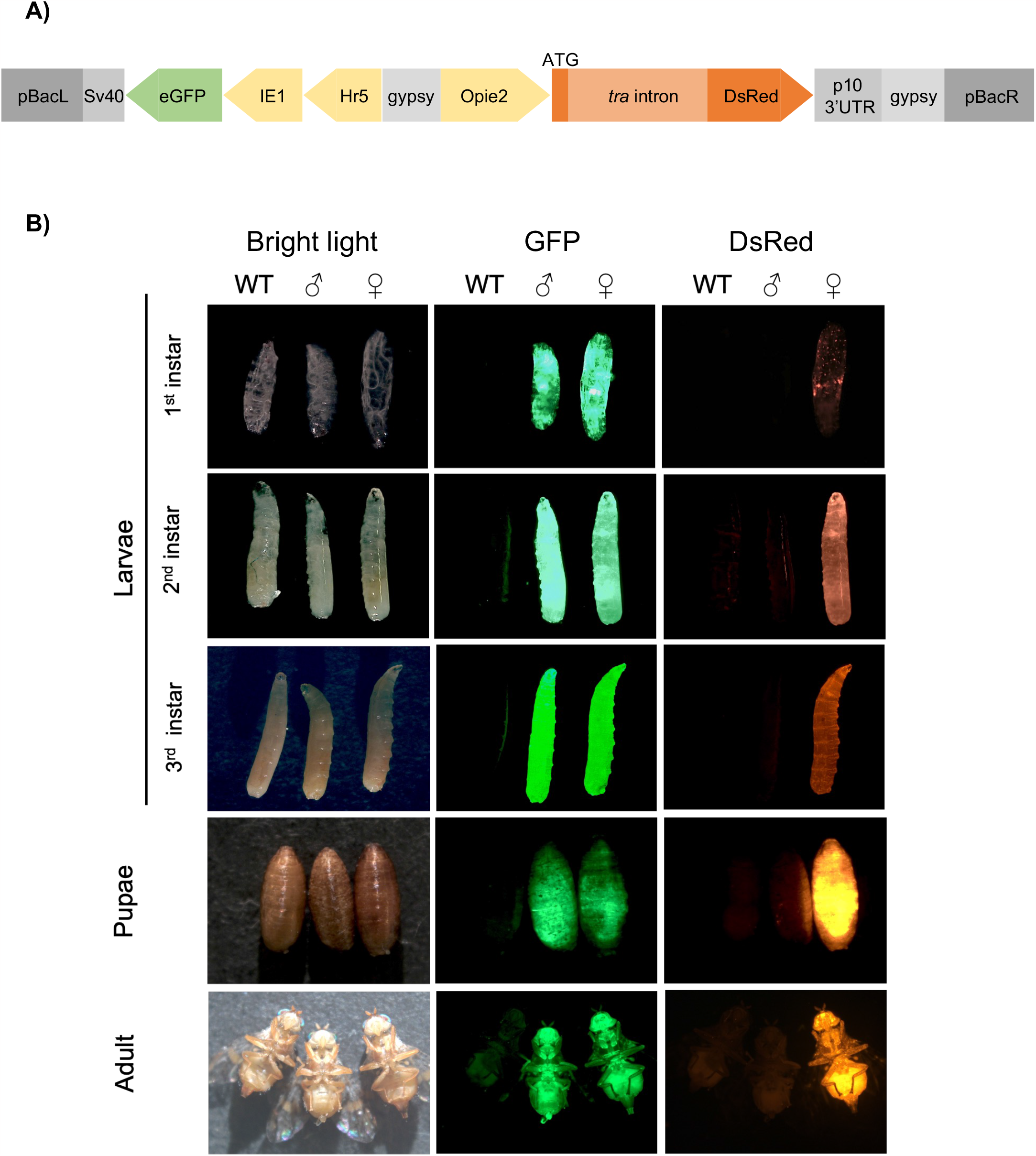
SEPARATOR system for sex-sorting of *Ceratitis capitata*. (A) A schematic representation of the Sexing Element Produced by Alternative RNA-splicing of Transgenic Observable reporter (SEPARATOR) cassettes, 795H1 and 795K1. Both cassettes contain two functional elements: constituent *Hr5IE1-eGFP-SV40* and *Opie2-DsRed-p10*, in which the DsRed coding sequence is separated by a *transformer* (*tra*) intron. The *tra* intron in 795H1 and 795K1 originates from *Ceratitis capitata* and *Anastrepha ludens*, respectively. (B) In the SEPARATOR strains the males produce eGFP only, whilst females express both eGFP and DsRed. The representative images of larval, pupal and adult stages of wild-type (WT), and homozygous male and female individuals are compared against each other under bright light, and green fluorescence protein (GFP), and red fluorescence protein (RFP) filters. All images were taken for the H-002 strain harboring the 795H1 construct. Images for all life stages were taken under bright light, and green fluorescence protein (GFP), and red fluorescence protein (RFP) filters.

**Figure 2:**
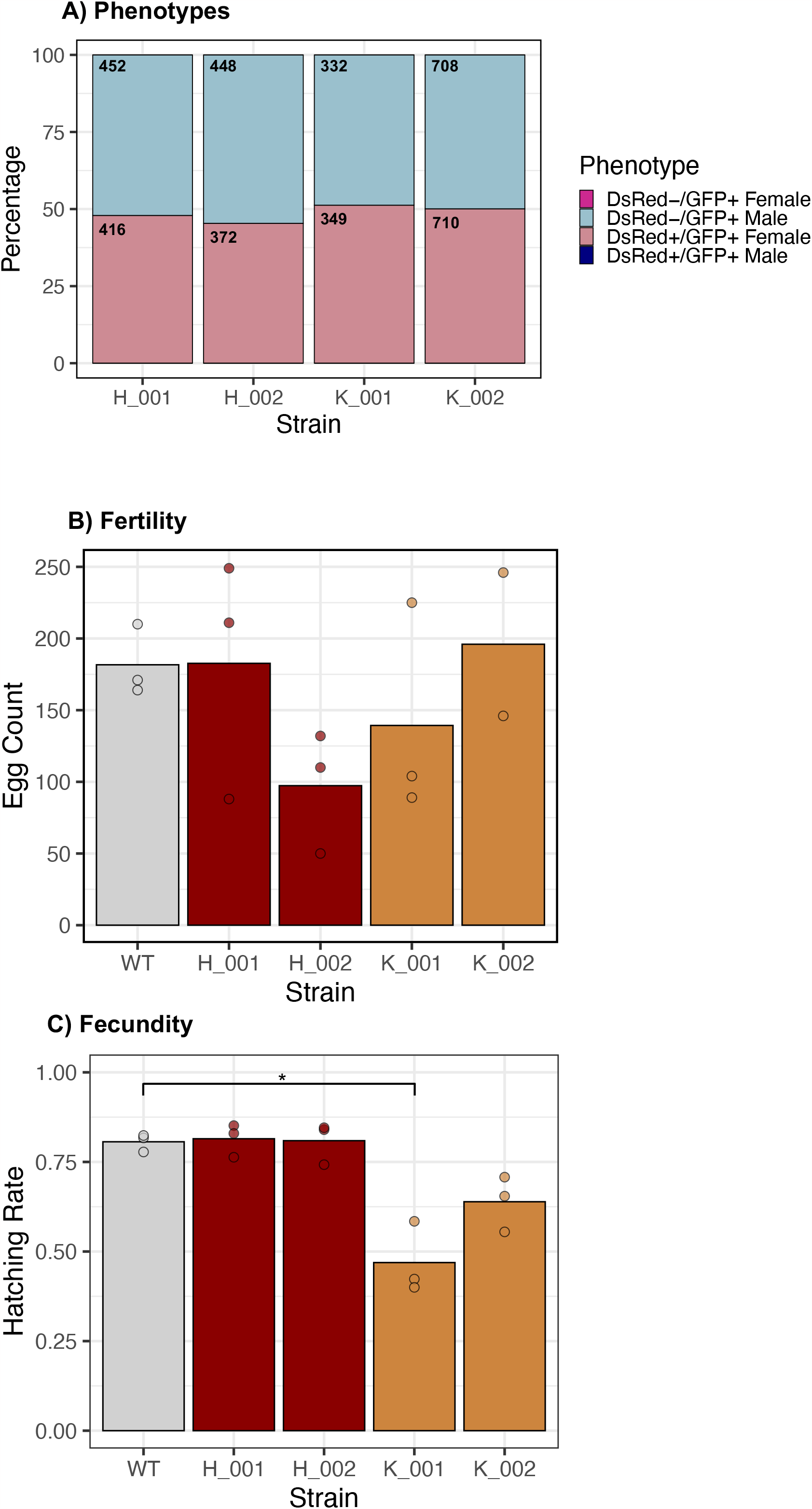
Characterization of transgenic SEPARATOR strains. (A) A stack graph showing the fluorescence phenotype distributions by sex of the four homozygous transgenic strains at the 9^th^ and 10^th^ generations with over 3,500 adult flies screened. Only the desired DsRed+/GFP+ females and DsRed-/GFP+ males were observed, whilst no DsRed+/GFP+ males or DsRed-/GFP+ females were observed. (B) Egg laying rates within strain crosses for all four strains was compared to wild-type through the egg laying rates within a 5-hour period. (C) Egg hatching rate within strain crosses for all four strains was compared to wild-type through the hatching rates of eggs laid within a 5-hour period. (A -C) H-001 and H-002 strains have the 795H1 cassette harboring the endogenous *Ceratitis capitata transformer* (*tra*) intron, while K-001 and K-002 carry the 795K1 cassette with the *Anastrepha ludens tra* intron. (A) Chi-squared tests showed no statistical significance in sex ratio distortion for any of the four strains. (B, C) Dunn test statistical significance was established at follows: p < 0.05 = * and the wild-type -transgenic strain significance is displayed on the graphs. (B, C) The bar levels represent the mean value whilst the dots represent raw values of the replicates. SEPARATOR stands for Sexing Element Produced by Alternative RNA-splicing of a Transgenic Observable reporter. (A -C) were constructed in RStudio.

### Fitness of fluorescent SEPARATOR strains

To assess whether the SEPARATOR cassettes are associated with notable fitness costs, we assessed the rates of egg laying and egg hatching rate for the four homozygous strains (Supplementary Table 4). Sibling crosses of homozygous individuals were conducted for each strain alongside wild-type controls. The egg laying rates were variable across wild-type and transgenic strains, although no statistically significant differences between wild-type and transgenic lines were determined through Kruskal-Wallis test and Dunn’s multiple comparison test (Figure 2b). It is of note, however, that there was a notable reduction in H-002 strain egg production. Meanwhile, the egg hatching rate of the 795H1-harboring strains with the endogenous *tra* intron was similar to wild-type (Figure 2c). The exogenous intron-containing 795K1 strains had reduced egg hatching rate compared to non-transgenic flies. Specifically, the egg hatching rate of K-001 was significantly reduced (p = 0.019), while the egg hatching rate of K-002 was reduced insignificantly (p = 0.072).

### Fluorescence patterns of SEPARATOR strains

We observed that the GFP fluorescence signal across the 4 homozygous strains was visibly similar. DsRed signal, however, was visually more intense in the females harboring the endogenous *C. capitata tra* intron (H-001 and H-002), compared to the females with the *A. ludens tra* intron (K-001 and K-002), irrespective of GFP fluorescence intensity in homozygous individuals (Supplementary Figure 2). We further investigated the patterns of fluorescence marker expression throughout the *C. capitata* life cycle (Figure 1b). We observed a clear distinction between males and females immediately upon hatching as larvae and increasingly throughout all later stages of the life cycle. Specifically, the signal was easily recognizable throughout the first, second and third instar larvae; pupae and adult development stages. At the late egg stage, GFP fluorescence is very clear in all individuals, whilst DsRed fluorescence was more difficult to differentiate (Supplementary Figure 3). This is because all eggs had a DsRed signal of variable intensity, which was stronger than in their wild-type counterparts.

We additionally carried out reverse-transcription PCR (RT-PCR) to amplify *Opie2-DsRed* for molecular verification of the sex-specific nature of *DsRed* splicing. cDNA, synthesized from total RNA, and gDNA were used to validate this phenomenon from adult males and females harboring either 795H1 or 795K1 (Supplementary Figure 4). We observed gDNA-derived fragments of similar sizes in both sexes, and cDNA-derived fragments of different sizes in males and females. The female-specific cDNA-derived fragments for both *C. capitata* (795H1) and *A. ludens* (795K1) *tra* introns expectedly amplified as single shortest fragments. One of the two male-specific transcripts was detected for 795H1 males, while multiple bands were observed for 795K1 males. These were more challenging to distinguish for the 795K1-harboring males as there are 5 alternative male-specific transcripts of *tra* in *Anastrepha* due to the presence of 3 male-specific exons (32). We deduced that *DsRed* in both cassettes is spliced in a similar fashion to the *tra* gene whereby DsRed translation is interrupted with male-specific stop codons. We also concluded that the *A. ludens tra* intron, despite incomplete sequence similarity, is functionally suitable to replace its endogenous counterpart in *C. capitata*.

## Discussion

In this study, we demonstrate a novel SEPARATOR approach to distinguish males from females in the global agricultural pest *C. capitata* with observed accuracy of 100%. This Y chromosome-independent and genetically stable strategy utilizes sex-specific splicing of *tra*, the master gene of sex-determination in Tephritids (29). We tested sex-specifically spliced *tra* introns from two species belonging to different genera within the *Tephritidae* family, the endogenous *C. capitata* and the exogenous *A. ludens. PiggyBac*-mediated transformations were used for line establishment in this study due to their high success rates across model and non-model insect species alike (46). To alleviate the insertion site-dependent effects on construct performance, we assessed 2 strains for each construct. In the system described above, females translate both a constitutive eGFP and the sex-specifically spliced DsRed fluorescence markers. The males, however, only produce eGFP as DsRed is not translated. The sex-specific splicing hypothesis is supported by the observed differential splicing patterns in males and females, altogether showing similar patterns of splicing to *tra* itself (Supplementary Figure 4). Both fluorescence signals are easily detectable at the early first instar larval stage and throughout all later stages of the *C. capitata* life cycle. Hence, there is an inherent suitability of this system to be implemented in larger-scale automated high-throughput sex separation. Furthermore, the system demonstrated in this study does not rely on antibiotics supply either for line maintenance (22, 25, 26) or for sex selection (24). Its cost efficiency, therefore, reinforces its scalability for SIT improvement purposes. Although the strains described here can be separated by sex at early development, the COPAS optical sorter system may be unsuitable for their separation as it requires water in which survival of *C. capitata* is immediately limited. Waterless optical sorter systems are therefore required to allow the most-efficient use of the SEPARATOR strains in SIT.

Importantly, here we observed that the *A. ludens tra* intron is capable of seamlessly mediating sex-specific splicing of *DsRed* in *C. capitata*. This highlights the conserved functional nature of *tra* and Tephritid sex determination altogether (27). When evaluating the fitness of transgenic lines, we noted high variation in egg laying among them, which may be explained by external stressors, integration site-specific effects and behavioral variation within the short experimental window. Additionally we noted a statistically significant reduction in one of the *A. ludens tra* intron-containing strains, whilst the endogenous intron-containing strains had similar egg hatching to wild-type controls (Figure 2c). This, in part, can be attributed to an additional burden of a larger exogenous intron, which may be alleviated when used in *A. ludens*. This is particularly useful in *A. ludens* as transgenesis in this species has a limited research history compared to *C. capitata* (26, 47, 48). The described system is also suitable for other cross-species applications due to the conserved function of *tra* throughout the wider Tephritidae family (32, 49, 50).

Remarkably, we observed no instances of DsRed-/GFP+ females, although this could be expected due to mutations in DsRed. On a mass rearing scale, however, this may become more frequent, hence a positive-selection system can be explored next whereby double-positive males are selected upon screening. This could employ pre-stop codon male-specific exons of *tra* or, alternatively, other genes spliced in a sex-specific manner such as *dsx* (51). Such positive selection system will mitigate risks of accidental female releases due to marker mutations. Additionally, the SEPARATOR system described here can serve as a backbone for novel strategies such as sex-specific expression of traditionally used selective markers. Specifically, this system can employ pupal coloration-involved genes such as *white pupae* and *ebony* can be targeted, which are both implemented in currently used Y-linked GSSs (15, 18). This can be achieved through site-specific integration of the *tra* intron into the coding region of the gene via homology-directed repair. This is particularly compelling for the use in the medfly as a protocol for such integration has just recently been used for gene drive establishment in the species (45).

## Methods

### Plasmid design and construction

Both 795H1 and 795K1 were constructed via Gibson assembly as described previously (43) into an existing *piggyBac* plasmid containing *Opie2* expressing DsRed and *Hr5Ie1* expressing eGFP. This plasmid was linearized using AvrII and BamHI restriction endonucleases. The *tra* intron of either *C. capitata* or *A. ludens* was PCR amplified from the corresponding genomic DNA preps and then inserted into the DsRed coding sequence immediately downstream of an ATG start codon (Supplementary Table 5). Complete sequence maps and plasmids are deposited at Addgene.org (#205482 for 795H1 and #205485 for 795K1).

### *C. capitata* rearing

The wild-type Benakeion strain of *C. capitata* used in this study was obtained from the Saccone Lab (University of Naples “Federico II”). Adult food consisted of a mixture of yeast and glucose in equal proportions and larvae were maintained using a carrot-based diet (52). Flies were consistently kept in 12:12 hour light: dark daily cycle at 26°C and relative humidity of 65%.

### *C. capitata* germline transformation

Plasmid microinjections were carried out into wild-type Benakeion strain embryos as previously described (53). The donor 795H1 and 795K1 *piggyBac* plasmids (500 ng/ml) were microinjected in combination with a helper plasmid encoding the ^*i*^*hyPBase* transposase (300 ng/ml) (54). The hatched G_0_ larvae were manually transferred onto larval diet and the surviving adults were crossed to virgin wild-type flies in a reciprocal manner. Fluorescence microscopy was used to identify marker-positive G_1_ progeny at the pupal stage of development, and selected adults were individually crossed with wild-type flies. Once established, homozygous transgenic lines were maintained through sibling crosses of 10 males and 20 females.

### Inverse PCR

Unique *piggyBac* construct integrations were analyzed through inverse PCR in selected marker positive G_1_ individuals. In brief, genomic DNA (gDNA) was first extracted via a modified protocol developed previously (55). The inverse PCR protocol was adapted from an established protocol (56) utilizing Sau3AI (New England Biolabs®) and HinP1I (New England Biolabs®) restriction endonucleases for initial gDNA digestion. *piggyBac*-specific primers from (57) were used for sequential PCR amplification. Sanger sequencing was carried out using Genewiz Inc. services and the resulting sequences were analyzed using the latest *C. capitata* genome reassembly (GenBank GCA_905071925.1).

### DsRed splicing confirmation

Adult males and females from transgenic lines were separately collected in TRIzol reagent (Ambion). RNA was extracted using an adjusted protocol from (58) and Maxima H Minus First Strand cDNA Synthesis Kit with dsDNase (ThermoFisher) was used for cDNA synthesis. In parallel, gDNA was extracted as described above. PCR was performed using Phusion High-Fidelity PCR Master Mix with HF Buffer (New England Biolabs®) or RedTaq DNA Polymerase 2X Master Mix (VWR Life Science) using primers designed in Geneious Prime 2023.1.2 (Supplementary Table 5). The amplicons were visualized using a 1% agarose gel.

### Sex sorting assay

For all homozygous strains two consecutive generations, G_9_ and G_10_, were screened to confirm system efficiency. Parental crosses between 10 male and 20 female homozygous individuals were established and eggs were collected twice 3 days apart for each cross. The offspring were reared until adulthood under regular conditions. At adulthood, all flies were assessed by 3 phenotypic parameters. These included phenotypic sex characteristics (male or female), as well as fluorescence marker phenotypes determined via separate screening with the GFP (GFP+ or GFP-) and RFP (DsRed+ or DsRed-) filters. MVX-ZB10 Macro Zoom Fluorescence Microscope System (Olympus) was used for all imaging and fluorescence screening.

### Fitness assays

To assess fitness costs of two copies of sex-sorting genetic cassette, the number of eggs laid and rate of egg hatching of homozygous cassette-carrying and wild-type flies were evaluated. Genetic crosses between 15 males and 25 females from each strain (H-001, H-002, K-001, K-002) were set up in biological triplicates. Simultaneously, crosses of 15 male and 25 female wild-type Benakeion strain adults were also established in triplicates. After 5 days, eggs oviposited within a 5-hour window were placed onto black filter paper on top of the larval diet and counted using ImageJ. The hatching rate was established 4 days after initial egg collection similarly through counting the remaining unhatched eggs.

### Figure generation and statistical analysis

All graphs were generated in RStudio using the ggplot() function. Statistical analyses including chi-squared, Kruskal-Wallis and Dunn tests were similarly performed in RStudio. Diagrams, gel and image-centered figures were constructed in Microsoft PowerPoint.

## Figure Legends

**Supplementary Figure 1:**
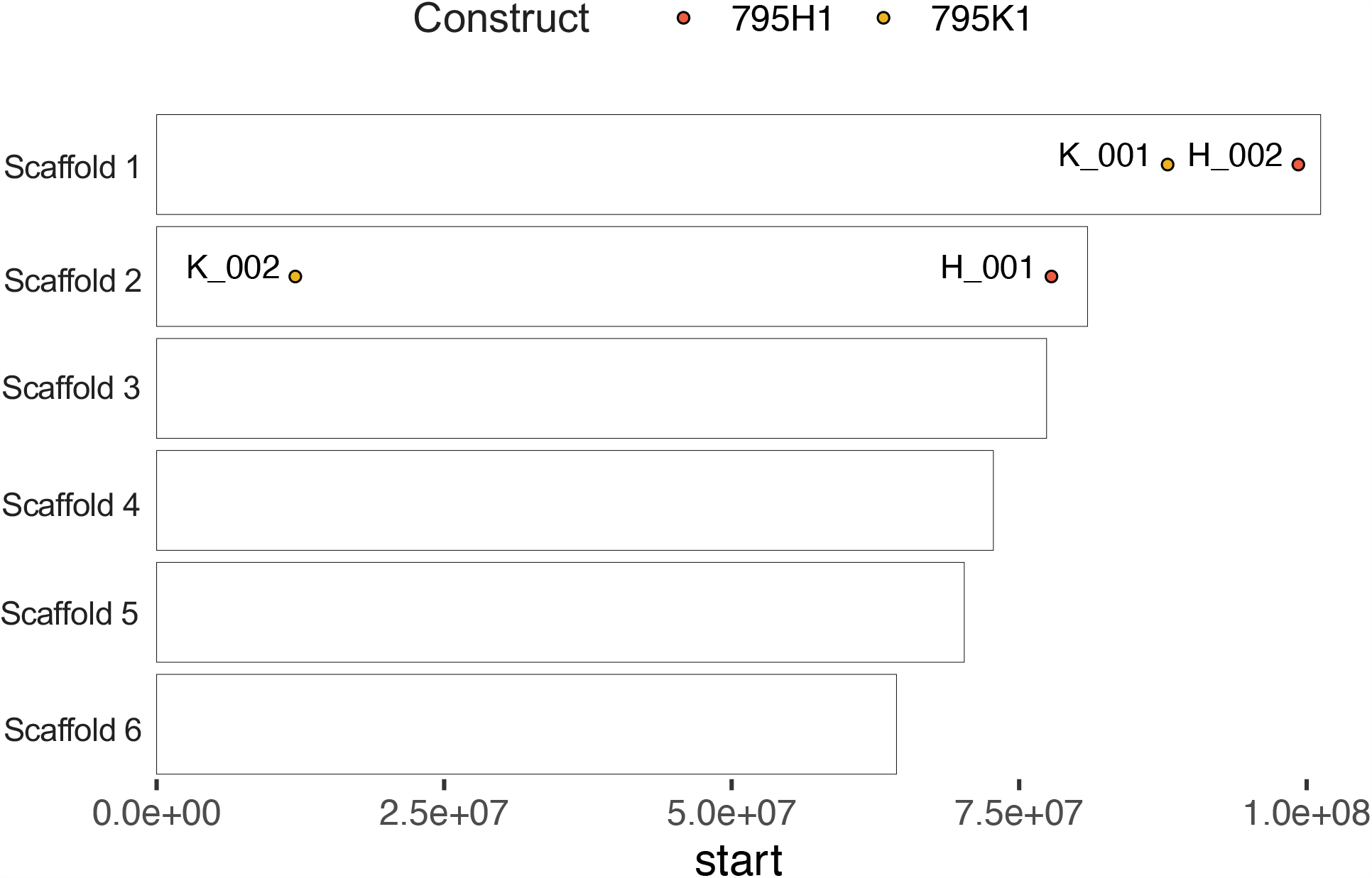
Inverse PCR integrations. The map of integrations of the 4 unique strains for the 795H1 and 795K1 constructs. H-001 and H-002 strains harbor the 795H1 cassette, while K-001 and K-002 strains harbor the 795K1 cassette. The figure was constructed in RStudio.

**Supplementary Figure 2:**
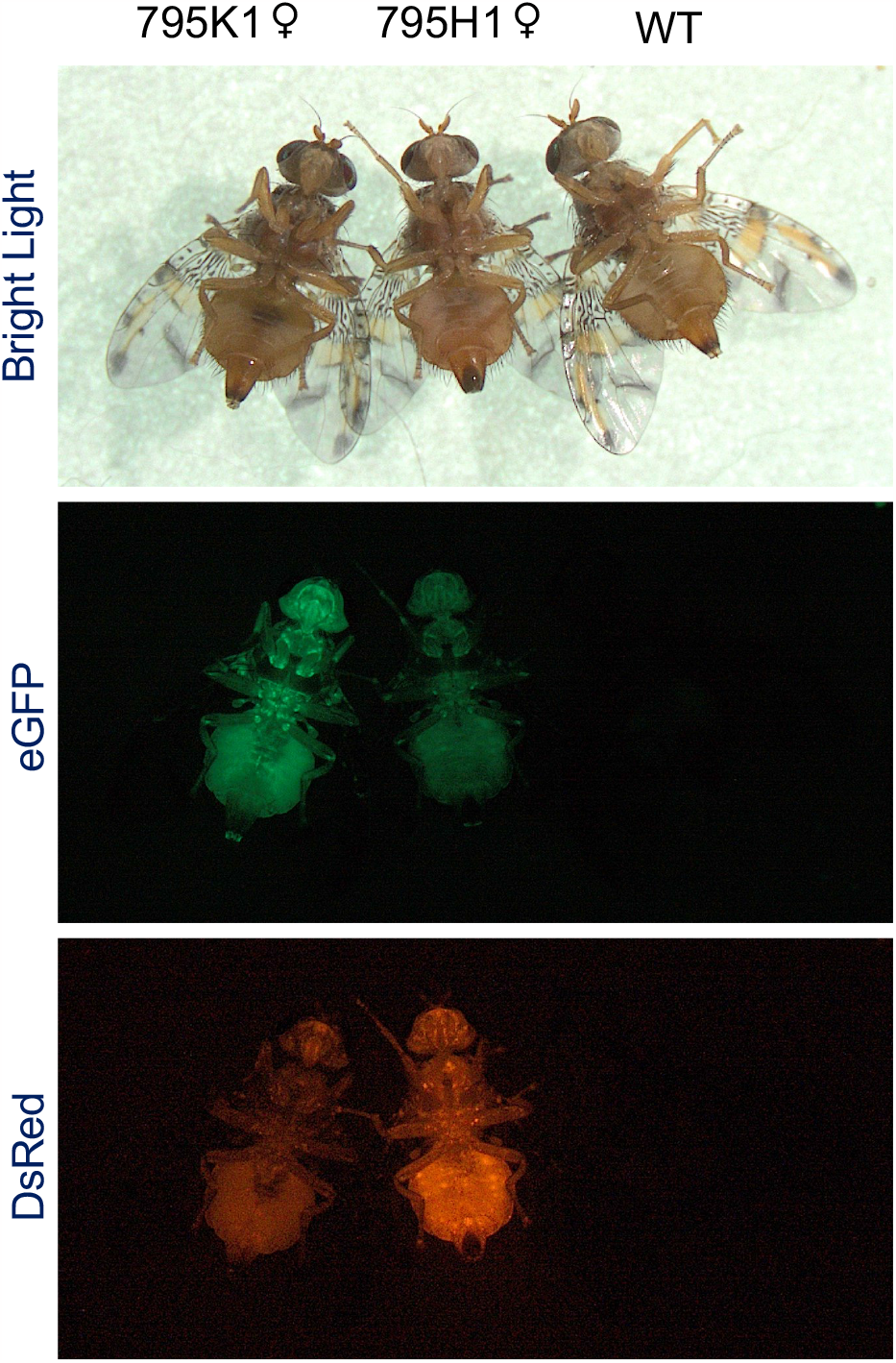
Images of 795H1 and 795K1 homozygous females. The homozygous females harboring the *Anastrepha ludens transformer* (*tra*) intron-containing 795K1 cassette have a weaker DsRed signal (left) compared to homozygous females harboring the *Ceratitis capitata* tra intron containing 795H1 cassette (middle). These were imaged alongside a wild-type female (right).

**Supplementary Figure 3:**
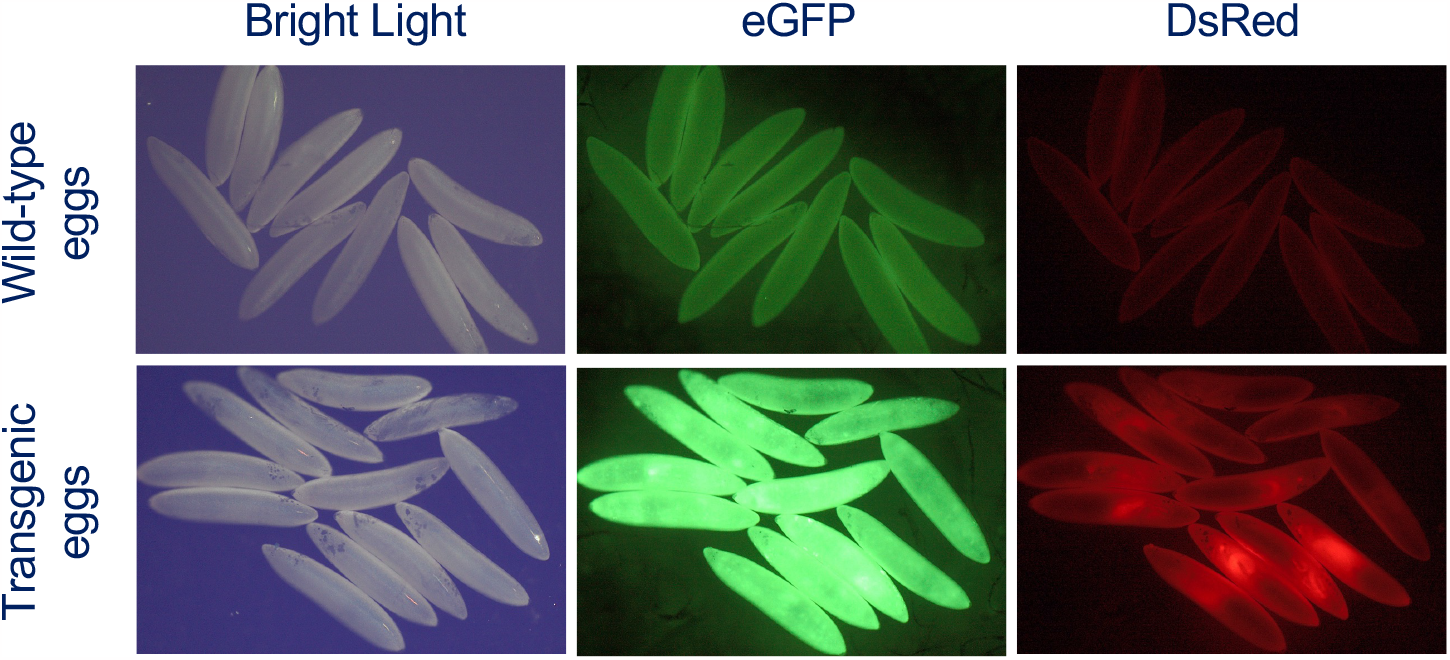
Images of transgenic and wild-type eggs. The egg images showcase the wild-type eggs, and the homozygous SEPARATOR eggs, all expressing GFP and a variable degree of DsRed.

**Supplementary Figure 4:**
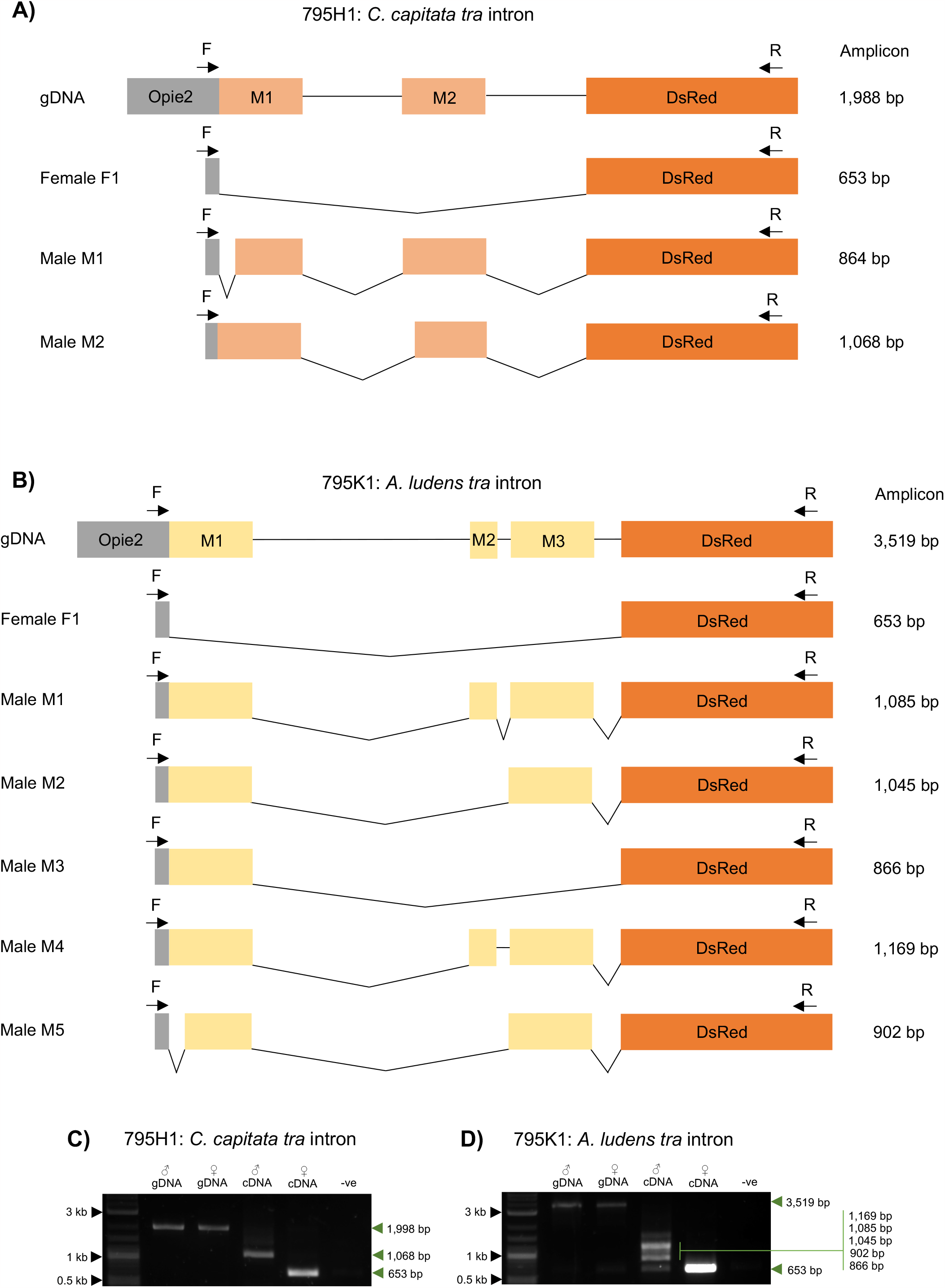
DsRed is spliced sex-specifically in all transgenic strains. Diagrams showcasing the expected sex-specific DsRed splicing patterns in (A) the 795H1-harboring and 795K1-harboring flies via the *transformer* (*tra*) intron from (A) *Ceratitis capitata* (29) and (B) *Anastrepha ludens* (32) accordingly. Forward and reverse primers (Supplementary Table 5), specific to the exogenous elements of both constructs, used in the PCR amplification are shown in (A) and (B) as F and R, respectively. (C) and (D) are annotated electrophoresis gel images with amplification from male and female genomic DNA (gDNA), and male, and female cDNA. A DNA ladder was run in the left-most lanes and negative controls in the right-most lanes.

**Supplementary Table 1:** G0 and G1 line raw data. Injection summary for 795H1 and 795K1 constructs. G0 data includes the number of injected eggs, hatched larvae and surviving adults. G1 data showcases the numbers of marker-positive adults screened for GFP and DsRed fluorescence markers.

| Transformation Construct | Eggs Injected | G0 Hatched Larvae | G0 Adults | G1 DsRed+ /GFP+ ♀ | G1 DsRed- /GFP+ ♂ |
| --- | --- | --- | --- | --- | --- |
| 795H1 | 250 | 210 | 84 | 31 | 11 |
| 795K1 | 250 | 141 | 52 | 4 | 2 |

**Supplementary Table 2:** Genomic integrations determined through inverse PCR. Inverse PCR outcomes, visualized in Supplementary Figure 2, obtained through mapping sequencing outputs to GenBank GCA_905071925.1 *C. capitata* genome re-assembly summarized by construct.

| Transformation Construct | Subline | Scaffold | Annotation | Integration Site |
| --- | --- | --- | --- | --- |
| 795H1 | H-001 | 2 | 77 806 718 | CACATCTCTAT**TTAA-PB-TTAA**CAGCTATAAT |
|  | H-002 | 1 | 99 273 433 | GTAAAACTTTT**TTAA-PB-TTAA**AACCAATTCA |
| 795K1 | K-001 | 1 | 87 907 601 | CCAGCAGTGAT**TTAA-PB-TTAA**CGTAGGTACG |
|  | K-002 | 2 | 12 057 725 | TTGGCAACTGT**TTAA-PB-TTAA**ACAGACATCA |

**Supplementary Table 3:** G9 and G10 line raw data. All adult flies in the G9 and G10 generations were phenotyped by sex and screened for GFP and DsRed fluorescence markers.

| Transformation Construct | Subline | DsRed+ /GFP+ ♀ | DsRed- /GFP+ ♂ |
| --- | --- | --- | --- |
| 795H1 | H-001 | 167 | 221 |
|  |  | 249 | 231 |
|  | H-002 | 156 | 219 |
|  |  | 216 | 229 |
| 795K1 | K-001 | 106 | 105 |
|  |  | 243 | 227 |
|  | K-002 | 330 | 307 |
|  |  | 380 | 401 |

**Supplementary Table 4:** Egg laying and egg hatching raw data. The unhatched eggs were counted via ImageJ twice, the first, immediately after collection, and then once more after 4 days. Number of hatched eggs was determined by calculating the difference between numbers of unhatched eggs in pictures 1 and 2.

| Construct | Strain | Replicate | Embryos | Hatched larvae |
| --- | --- | --- | --- | --- |
| N/A | WT | 1 | 171 | 133 |
|  |  | 2 | 164 | 134 |
|  |  | 3 | 210 | 173 |
| 795H1 | H-001 | 1 | 249 | 212 |
|  |  | 2 | 88 | 73 |
|  |  | 3 | 211 | 161 |
|  | H-002 | 1 | 50 | 42 |
|  |  | 2 | 132 | 98 |
|  |  | 3 | 110 | 93 |
| 795K1 | K-001 | 1 | 104 | 44 |
|  |  | 2 | 89 | 52 |
|  |  | 3 | 225 | 90 |
|  | K-002 | 1 | 146 | 81 |
|  |  | 2 | 253 | 179 |
|  |  | 3 | 246 | 161 |

**Supplementary Table 5:**
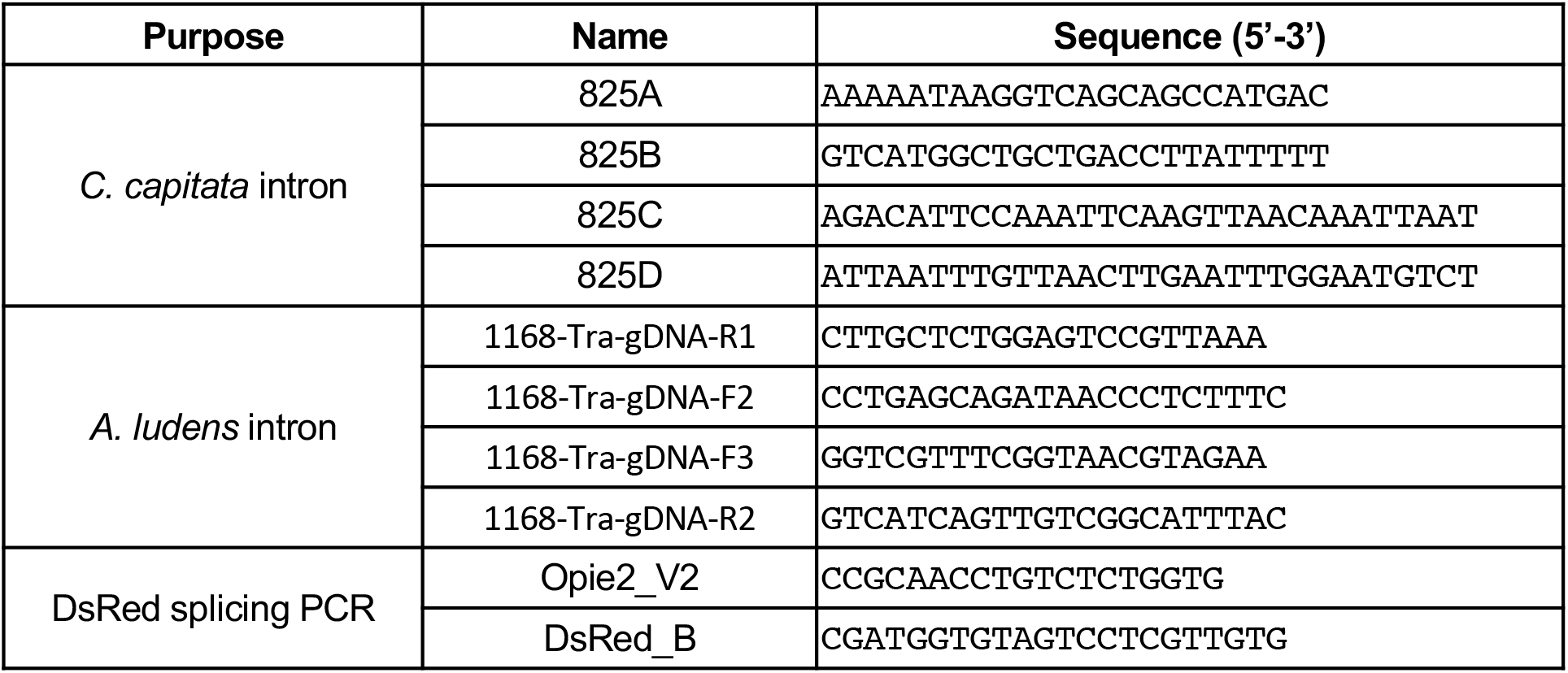
Primer summary. The primers used for cloning of 795H1 and 795K1 constructs, and the primers used for DsRed splicing confirmation (Supplementary Figure 3).

## Data availability

Complete plasmid sequences are available at addgene.org (#205482 and #205485). All data used to generate figures is provided in the Supplementary materials. Medfly transgenic lines are available upon request to A.M.

## Author contributions

O.S.A, N.P.K, A.M and E.B conceived the project and directed the research. J.L and N.P.K designed constructs and J.L performed the cloning. A.M and S.D designed the experiments and S.D performed the medfly and molecular validation experiments. A.M performed the germline transformation. A.M and S.D analyzed the data. All authors contributed to writing and editing of the manuscript and approved the final article.

